# Mapping of transgenic alleles in plants using a Nanopore-based sequencing strategy

**DOI:** 10.1101/529230

**Authors:** Shengjun Li, Shangang Jia, Lili Hou, Hanh Nguyen, Shirley Sato, David Holding, Edgar Cahoon, Chi Zhang, Tom Clemente, Bin Yu

## Abstract

Transgenic technology was developed to introduce transgenes into various organisms to validate gene function and add genetic variation for the development of beneficial input or output trait over 40 years ago. However, the identification of the transgene insertion position in the genome, while doable, can be cumbersome in the organisms with complex genomes. Here, we report a Nanopore-based sequencing method to rapidly map transgenic alleles in the soybean genome. This strategy is high-throughput, convenient, reliable, and cost-efficient. The transgenic allele mapping protocol outlined herein can be easily translated to other higher eukaryotes with complex genomes.

## Introduction

Transgenic technologies that introduce genetic variation into bacteria, animals and plants were developed in 1972 (Cohen *et al.*, 1972), 1974 (Jaenisch and Mintz, 1974) and 1982 (Barton *et al.*, 1983), respectively. They have become a valuable resource to enhance genetic variations and to gain insight of gene function. In higher plants, a single cope or multiple copies of transgenes are randomly inserted into the genome (Kim *et al.*, 2007; Weising *et al.*, 1988). Expression levels of a transgene are often influenced by the genomic context surrounding the transgenic allele and the complexity of the genome (Butaye *et al.*, 2004; Day *et al.*, 2000; van Leeuwen *et al.*, 2001; Weising *et al.*, 1988). Moreover, the transgene insertion position may also affect the function of surrounding genes (Azpiroz-Leehan and Feldmann, 1997; Weising *et al.*, 1988). Importantly, prior knowledge of map position of a transgenic allele is beneficial when breeding programs begin to introgress the allele into elite germplasms. Consequently, there is a need to efficiently and accurately characterize transgenic alleles in higher plants

Strategies have been developed for mapping of transgenic alleles (Guo *et al.*, 2016; Lepage *et al.*, 2013). Complexity of the transgenic locus can be estimated through multiple approaches including Southern blot analysis (Southern, 1975), quantitative PCR (Ingham *et al.*, 2001) and droplet PCR (Glowacka *et al.*, 2016). One of the first methods used to successfully map a transgenic allele in higher plants was plasmid rescue. This strategy involves restriction enzyme digestion of the host genome containing the transgenic allele, cloning the cleavage products into plasmid and selection of the plasmid containing the transgene fragment (Nan and Walbot, 2009). Subsequent methods for mapping transgenic alleles are also primarily PCR based, include Thermal Asymmetric Interlaced PCR (TAIL-PCR) (Liu and Chen, 2007; Liu *et al.*, 1995), Adaptor PCR which is sometimes referred to as anchored PCR (Singer and Burke, 2003; Thole *et al.*, 2009), and T-linker PCR that utilizes a specific T/A ligation (Yuanxin *et al.*, 2003). However, these methods are often challenging to scale-up for high-throughput (Guo *et al.*, 2016; Ji and Braam, 2010). Moreover, failure to map transgenes can happen due to the complexity of the transgenic locus and/or issues associated with the genomic context about the transgenic allele (Wahler *et al.*, 2013). The next-generation Illumina sequencing technology is a method that can map transgenic alleles in plants due to its depth of sequencing capacity (Guo *et al.*, 2016; Lepage *et al.*, 2013; Polko *et al.*, 2012). However, because this method produces short reads, a high degree of sequencing depth is needed, especially in crops that have large genomes that are rich in repetitive sequence. This in turn, impacts the cost per transgenic locus mapped. In addition, short-read sequencing data is challenging to resolve transgene insertion position in many plant species, such as soybean, due to issues related with genome rearrangements and copy-number variations, which may lead to inaccurate mapping locations.

Recently, single molecule real-time (SMRT) sequencing technologies have been developed that provide long-read sequencing datasets. These SMRT platforms developed by Pacific Biosciences (PacBio^®^) and Oxford Nanopore Technologies^®^ offer significant attributes for genotyping plant species. The most significant benefit is long read lengths, with Pacbio^®^ platform generating up to 60 kb reads, and Nanopore^®^ reads being up to ~ 1Mb (Jain *et al.*, 2018; Lu *et al.*, 2016). Both technologies have been used in genome assembly (Badouin *et al.*, 2017; Jain *et al.*, 2018; Michael *et al.*, 2018; Rhoads and Au, 2015; Schmidt *et al.*, 2017)). The MinION device, which was developed by Nanopore^®^ technology and entered the market in 2014, is a portable apparatus with less than 100g in weight. Furthermore, it is compatible with a PC or laptop with USB 3.0 ports (Jain *et al.*, 2016) making it a flexibility attribute permitting use outside of a laboratory setting (Castro-Wallace *et al.*, 2017). In addition, compared with PacBio^®^, the Nanopore Technology apparatus is affordable in most laboratories. Thus, the MinION platform provides potential for a high-throughput, cost-effective strategy to map transgenic alleles in plant species with complex genomes.

Described herein is a Nanopore Technology®-based platform pipeline designed for high-throughput mapping of transgenic alleles in plant species. Employing a target enrichment approach using a combination of oligo probes to capture DNA fragments containing the transgenic allele, permitted the rapid identification of map position of 51 transgenic alleles in a single 1D sequencing-run. The calculated cost incurred by the procedure to map 51 transgenic alleles is estimated to be $1,360, and the results are generated within one week. The reads with the transgenic allele averaged in the hundred, for each sample, suggesting that pooling can be further enlarged. These results demonstrate that this Nanopore^®^-based sequencing method is rapid, convenient, reliable, cost-efficient and high-throughput.

## Materials and Methods

### Soybean growth condition

The soybean plants were grown in controlled greenhouse condition with 14 hour photoperiod and 28/26°C day/night temperature. The soybean plants harboring the *Ds* element are in the Thorne genetic background.

### DNA extraction and shearing

DNA from of soybean leaves were extracted using CTAB method (Healey *et al.*, 2014) and purified with DNeasy Plant Mini Kit (69104, QIAGEN). 6 μg genomic DNA in a total of 150 μl nuclease free water was sheared into ~8 kb with g-TUBEs (520079, Covaris) by following manufacturer’s instruction.

### DNA barcodes and Enrichment of the *Ds* element-containing fragments

1 μg sheared DNA fragments were end-repaired with Ultra II End-prep enzyme mix (E7546L, NEB) for 5 minutes at 20°C and 5 minutes at 65°C using a thermal cycler, followed by purification with the AMPure XP beads in a 1.5 ml DNA LoBind Eppendorf tube. After end-repaired, DNA fragments were ligated to the Barcode Adapter from the barcode Kit 1D (EXP-PBC001, Nanopore) using Blunt/TA Ligase Master Mix (M0367L, NEB). Following purification with AMPure XP beads, the DNAs were ligated to the Barcode (EXP-PBC001, Nanopore) using LongAmp Taq (M0287S, NEB). The barcoded DNA library was then purified with AMPure XP beads. After barcoding, the library was purified with pheno/chloroform method, and diluted with 4.8ul H2O+8.5ul xGen 2X Hybridization buffer, then add 2.7ul xGen Hybridization enhancer (1072281, Integrated DNA Technologies, IDT) and 1ul probe. Then hybridization was performed at 65°C for 4h in a thermal cycler. After hybridization, the targets were captured by the Dynabeads M-270 Streptavidin beads (65-305, Thermo Fisher Scientific) that recognize the dualbiotinylated probe. After washing with Stringent Wash Buffer and Wash Buffer I, II, III by following the manufacture’s protocol, the captured target fragments were amplified for 12 cycles with primers recognizing the barcode using LongAmp Taq at the PCR condition: 15 seconds at 98°C, 30 seconds at 60°C, 6 minutes at 72°C. The resulting PCR products were purified with AMPure XP beads, which were subjected to second round enrichment (step 3 and 4), or library construction following manufacturer’s instruction. The 5’ dual biotinylated probe was synthesized from IDT and its sequence is shown probe in Supplementary Table S1.

### Library Construction and Sequencing

Following target enrichment, barcoded libraries were pooled and 1 μg samples were end-repaired with the Ultra II End-prep enzyme, purified with the AMPure XP beads and then ligated to the sequencing adaptor (SQK-LSK108, Nanopore) with the Blunt/TA Ligation Master Mix. After purification with the AMPure beads, the adapted DNA libraries were sequenced in the flow cells (R9.4 version, FLC-MIN106, Nanopore). After 20-24 hours, the sequencing was stopped.

### Assessment of target enrichment efficiency

To assess the target enrichment, 2% of samples were used as templates to perform quantitative PCR (qPCR) using SYBR Green PCR Master Mix (Bio-Rad) with primers recognizing the Ds element or an unrelated intergenic region in soybean chromosome 7. The primer sequences are shown in Supplementary Table S1.

### PCR validation

PCR reaction was performed with primers listed in Supplementary Table S1 using the condition: 95 °C 2 min; 95 °C 30 sec, 50 °C 30 sec, 72 °C 1:20 min for 34 cycles; 72 °C 5 min. The PCR products were isolated with 1% agarose gel and visualized by Ethidium bromide staining.

### Bioinformatics analysis

All barcoded reads were de-multiplexed and adapters were trimmed off using the Porechop version 0.2.1 (https://github.com/rrwick/Porechop) with default parameters. To identify reads with the *Ds* target sequence, the *Ds* target sequence was searched against trimmed reads for each sample with E-value ≤ 10^−3^. For all hits with the *Ds* target sequence, the 5’ end and 3’ end sequences of the *Ds* target sequence were scanned on each read to identify long reads with one or two complete ends of the *Ds* target sequence. Sequences on 5’ end and/or 3’ end sequences of long reads beyond the *Ds* target sequence, if length > 20bp, were recorded as flanking sequences, which come from soybean genome. The flanking sequences were undergone blast searches against the soybean genome (v1.0). Uniquely aligned hits with aligned length > 200bp and > 80% sequence identity were kept. The genomic location for each flanking sequence were determined based on its alignment. The insertion sites were determined based on statistically enriched flanking sequences. The zero-inflated Poisson regression was used to model count data that has an excess of zero counts. All read counts were fitted into the Zero-inflated Poisson regression model with the R package, ZIM. For each peak of read counts, to determine if it was a significant peak, a P-value was calculated as the probability of observing a count value equally as extreme, or more extreme, than the given read count based on the fitted distribution

## Results

### Mapping of maize *Ds* transpositions in the soybean genome through MinION sequencing without target enrichment

To evaluate the potential application of MinION sequencing to map transgenic alleles, a soybean line, which contains a transgene stack harboring the maize Activator (*Ac*)/Dissociation (*Ds*) transposon system were used. The *Ac* transposase is controlled by the 35S CaMV promoter, and the *Ds* element harbors the cassava vein mosaic virus promoter (CsVMV) as an activation tag. The selected soybean lines were previously genotyped via Southern blot analysis to ascertain the presence of the *Ds* loci and the absence of *Ac* allele, along with mapping of the Ds allele using TAIL-PCR (Fig. 1A and Supplementary Fig. S1). To assess the power of MinION sequencing to map transgenic alleles, genomic DNA isolated from one of the selected genotyped soybean lines carrying the *Ds*-activation tag was sequenced on the FLO-MIN106 flow cell following the 1D sequencing protocol without DNA fragmentation (Fig. 1B). A 24-hour sequencing run produced approximately one million reads, resulting in about 2.8 Gb of sequence data (Table 1). Mining the sequence data for *Ds* element revealed two reads containing the *Ds* element (Table 1). One read was 957 bp covering partial *Ds* element flanked by 370 bp sequence at 3’ end, and the other was 6,806 bp, containing the full-length *Ds* element flanked by 2,347 bp 5’ upstream sequences and 3,047 bp downstream flanking the Ds sequence (Fig. 1C). The identified *Ds* junction fragment sequences were mapped to the soybean Glyma.15g128600 gene (Fig. 1C), in agreement to the TAIL-PCR results. To further validate the sequencing and TAIL-PCR outcomes, PCR reactions were carried out with a primer set designed to span the *Ds*/junction about the insertion site (Fig. 1A). The data revealed a 360 bp PCR product amplified from the endogenous Glyma.15g128600 gene when control DNAs were used as templates, and a 1526 bp fragment predicted to carry the *Ds*/junction target sequence amplified from DNAs of the transgenic soybean plants (Fig. 1D). These results demonstrate the potential of MinION sequencing to map a transgenic allele in the soybean genome. However, given the few reads that contain the *Ds,* refinement in the genomic DNA processing steps would be required for a high throughput/cost effective mapping pipeline with this technology.

**Table 1.** Sequencing result of one line without enrichment.

|  | Total reads<br>number | Longest read<br>(bp) | Target reads<br>number | Percent of<br>target reads | Longest read<br>with targets<br>(bp) |
| --- | --- | --- | --- | --- | --- |
| Line 1 | 1061117 | 351899 | 2 | 0.00019 | 6806 |

Note: Longest read indicates the longest read in all readings.

**Fig. 1.**
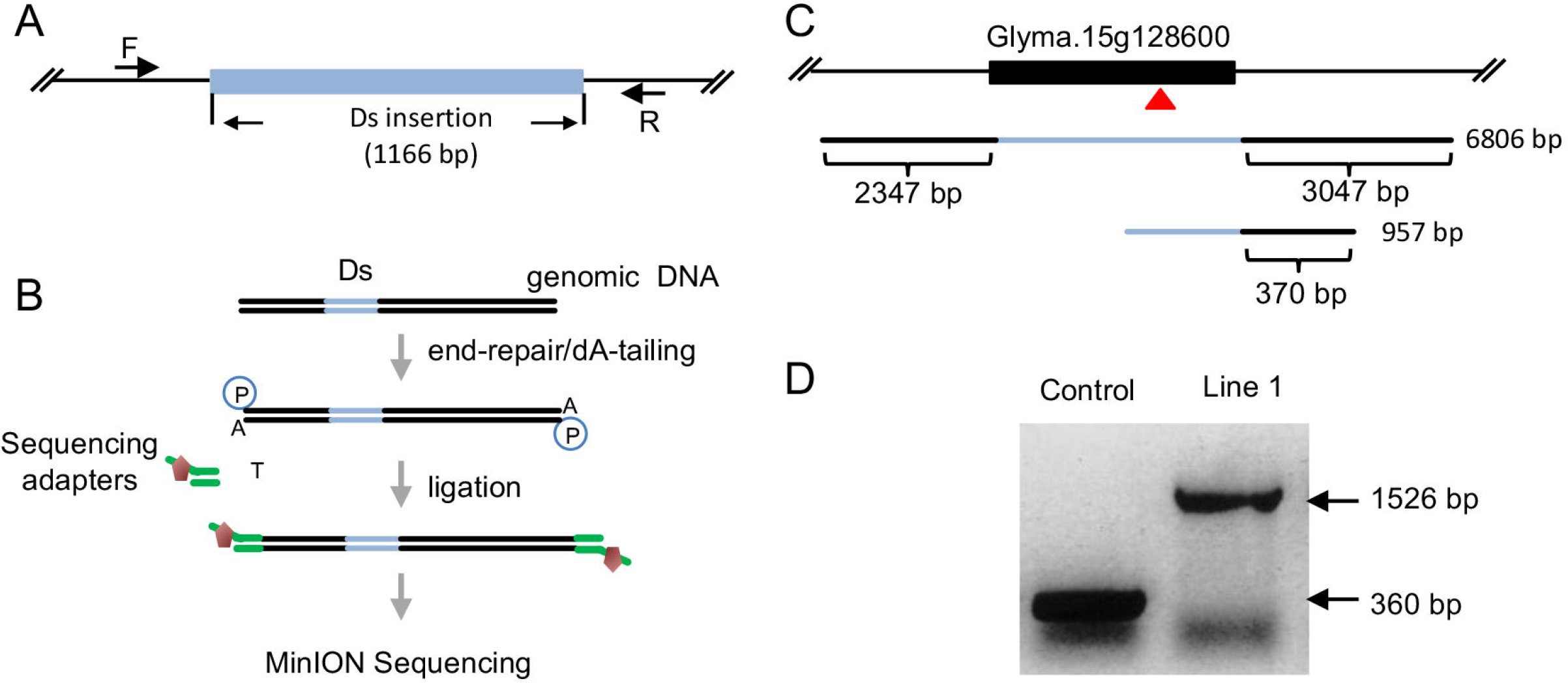
MinION sequencing without *Ds*-enrichment. **(A)**The schematic diagram of *Ds* insertion in soybean genome. The length of *Ds* insertion is 1166bp. The positions of forward (F) and reverse (R) primers used for PCR genotyping are shown. **(B)**Workflow of direct genome sequencing without target-enrichment. Genomic DNA was end-repaired and dA-tailed, ligated with sequencing adapters and sequenced on the FLO-MIN106 flow cell. **(C)**The schematic diagram of the *Ds* insertion in Glyma.15g128600 gene. Two reads are shown. The first one covers 2347bp in the 5’ flanking region and 3047bp in the 3’ flanking region. The second one contains 370bp flanking sequence in the 3’ region. **(D)**PCR validation of the *Ds* insertion in Line 1. Thorne was used as control plant. The length of DNA fragment without the *Ds* element in control plant is 360 bp, while the fragment length from Ds-containing Line 1 is 1526 bp.

### Target enrichment of transgenic allele to improve mapping throughput with MinION sequencing

To improve read counts around the junction of a transgenic allele, a PCR-based method to enrich the target sequences in the DNA library (Fig. 2) was developed. To test the enrichment protocol, DNA from two soybean lines carrying a *Ds* activation tag allele, previously characterized via Southern blot and mapped by TAIL-PCR, were used. The enrichment protocol incorporated steps for fragmentation of DNA to approximate 8 kb, end-repairing and dT-tailing, with subsequent ligation to barcode adapters and PCR-barcoding (Fig. 2). The resultant reaction products were subjected to a 120nt 5’-dual biotinylated probe designed to capture the transgenic *Ds* allele (Fig. 2). Following the probe capture step, the probe-captured fraction was re-amplified by PCR and products were pooled for sequencing (Fig. 2). Total readings obtained were 357765 and 326189 for Line 2 and Line 3, respectively (Table 2). The average read length of Line 2 was 2426 bp with the longest read of 20453 bp (Table 2), while the average read length of Line 3 was 2445 bp with the longest read of 48971 bp (Table 2). Among the reads obtained implementing the enrichment steps, 203 and 438 contained the *Ds*-allele sequence, for lines 2 and 3, respectively, which correctly mapped to gene calls, Glyma.19g105100 and Glyma.11g247400, respectively (Fig. 3A, 3B, and Table 2). The map positions were re-confirmed using PCR analyses incorporating a primer set designed to amplify *Ds*/junction fragment region (Fig. 3C).

**Table 2.** Sequencing result of two lines with one-round enrichment.

|  | Total reads<br>number | Longest read<br>(bp) | Target reads<br>number | Percent of<br>target reads | Longest read<br>with targets<br>(bp) |
| --- | --- | --- | --- | --- | --- |
| Line 2 | 357765 | 20453 | 203 | 0.057 | 6524 |
| Line 3 | 326189 | 48971 | 438 | 0.134 | 6725 |

Note: Longest reads indicate the longest reads in the individual barcoded lines.

**Fig. 2.**
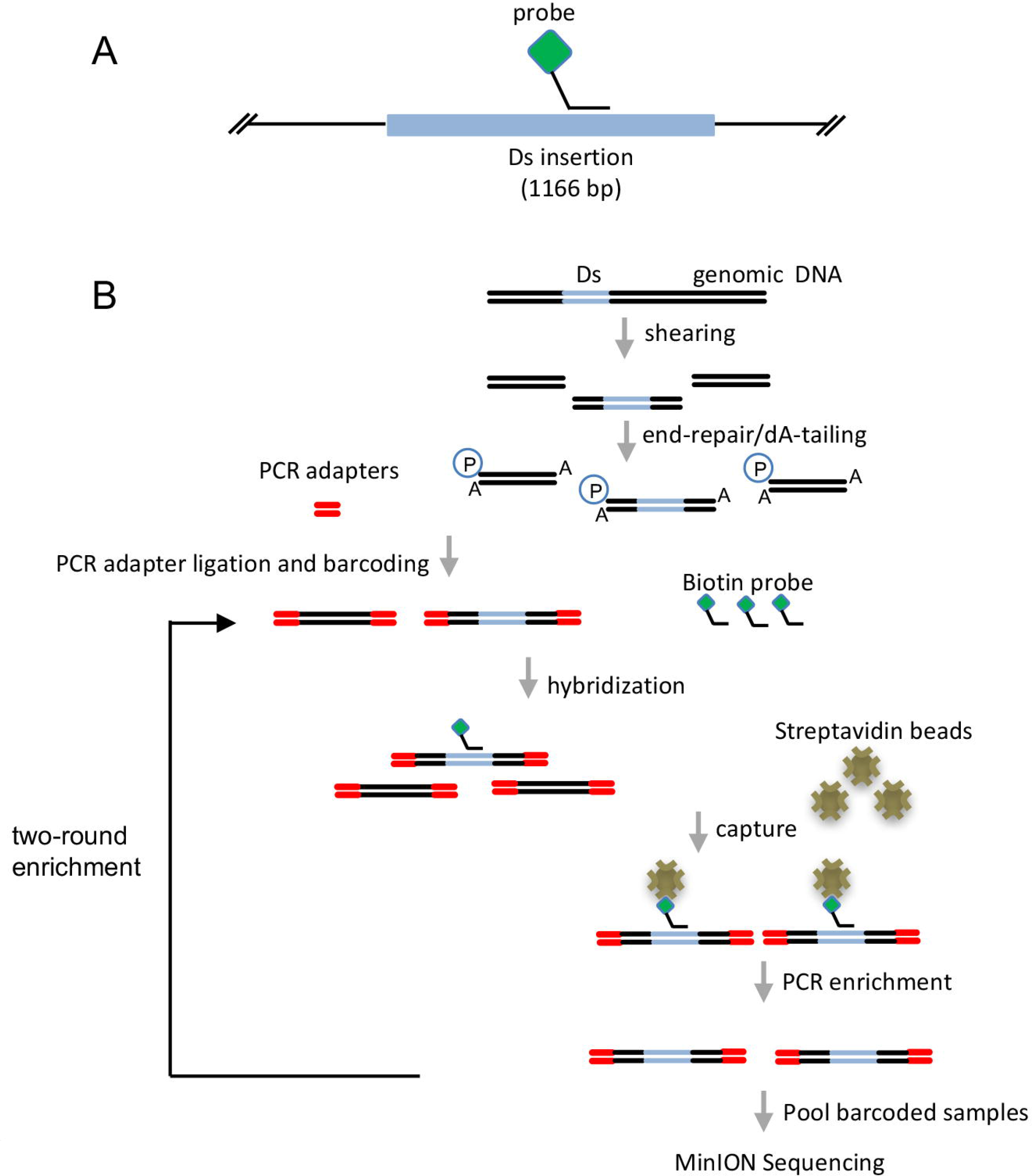
The workflow of the enrichment of *Ds*-containing fragments in DNA libraries. **(A)**The schematic diagram of oligo probe used to capture the *Ds* element. The probe is dual biotinylated at 5’ end (green diamond). **(B)**The workflow of sequencing the enriched *Ds*-containing DNA fragments. Genomic DNA was sheared and ligated to PCR barcode adapters. The *Ds*-containing fragments were enriched one or two rounds. The enriched fragments were pooled and sequenced.

**Fig. 3.**
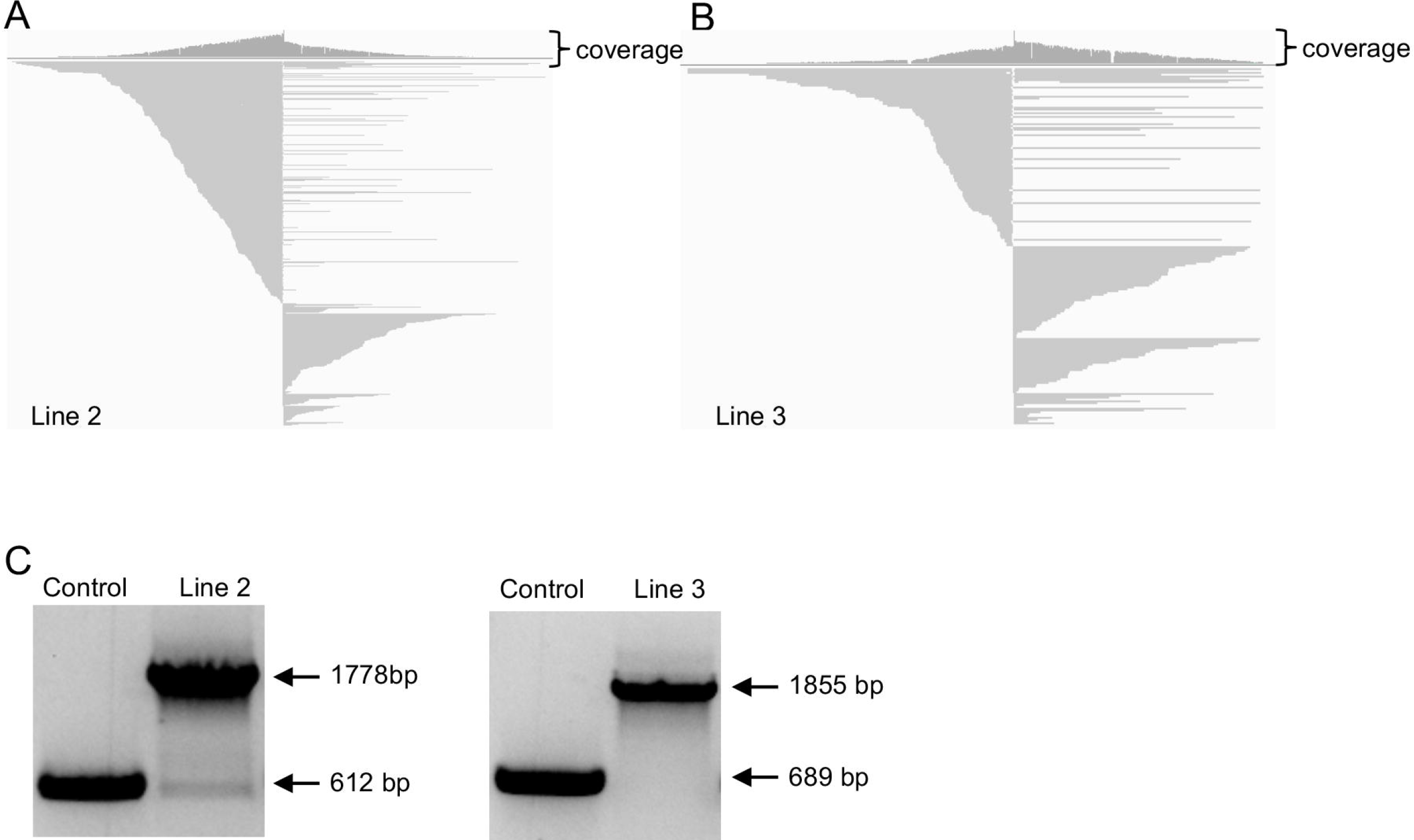
Sequencing results after one-round enrichment of the *Ds*-containing fragments. **(A)** and **(B)**Schematic diagram of the flanking sequences of Line 2 (A) and Line 3 (B). Partial sequences of reads were shown. **(C)**PCR validation of the Ds insertion in Line 2 and Line 3. Thorne was used as control plant. The lengths of the DNA fragment without the *Ds* element are 612bp for Line 2 and 689 bp for Line 3. With the *Ds* elements, the lengths of the DNA fragments are 1778 bp for Line 2 and 1855 bp for Line 3.

Given the high number of reads containing the *Ds* element, following the targeted enrichment approach, the method appeared to be amendable for higher throughput by increasing sample pool size. To this end, 15 soybean lines previously ascertained to harbor a single *Ds* element (Line 4-Line 18) were selected for integrating a pooling strategy with the targeted enrichment method. Here five DNA pools, each of which contained DNAs from three soybean lines (Table 3 and Supplementary Fig. S2), were prepared. Following the first target enrichment step, the pools were subjected to an additional round of purification to increase coverage of the *Ds*-containing DNA fragments (Fig. 2). Subsequent to each purification step, a quantitative PCR (qPCR) was used to estimate the relative enrichment level of target fragment compared with an unrelated DNA region that served as an internal control. After one round of enrichment, the ratio of target fragments to the unrelated region was enriched 132-1120x across all DNA pools (Fig. 4A). Following two rounds of purification the enrichment ratio ranged from 7469 to 238193 times in the pools (Fig. 4B). MinION sequencing of the double enriched products resulted in total number of reads ranged from 117266 to 523192 across the pools (Table 3), with reads containing the *Ds* sequence ranging from 1856 to 36388 in the pools (Table 3). These results were translated to ratios of reads containing the target sequence per total read counts for each pool in the range of 0.53 to 6.95% (Table 3). The average length of these Ds-containing reads was longer than 2Kb and the majority (>99%) of these readings were longer than 1.2 Kb (Fig. S3). The *Ds*-containing reads of each DNA pool were successfully mapped to three positions in the soybean genome (Supplementary Table S2; Fig. 4C showing a position of readings at soybean genome from Pool 4), reflecting that the pools each contained three independently integrated *Ds* elements within the soybean genome. The predicted mapped locations identified in Pool 4 were subsequently verified by PCR using primer sets that spanned the *Ds* element/soybean genome junction (Line 13-Line 15; Fig. 4D).

**Table 3.** Sequencing result of the 15-sample pools.

|  | Line<br>number | Total reads<br>number | Longest read<br>(bp) | Target<br>reads<br>number | Percent of<br>target reads | Longest<br>read with<br>targets (bp) |
| --- | --- | --- | --- | --- | --- | --- |
| DNA pool 1 | Line 4-6 | 351722 | 16352 | 1856 | 0.53 | 5100 |
| DNA pool 2 | Line 7-9 | 490852 | 21457 | 30937 | 6.30 | 9317 |
| DNA pool 3 | Line 10-12 | 117266 | 8000 | 3165 | 2.70 | 5770 |
| DNA pool 4 | Line 13-15 | 523192 | 25983 | 36388 | 6.95 | 13213 |
| DNA pool 5 | Line 16-18 | 234809 | 14215 | 5412 | 2.30 | 6008 |

Note: Line number indicates the pooled Ds-containing lines. Longest reads indicate the longest reads in each pool.

**Fig. 4.**
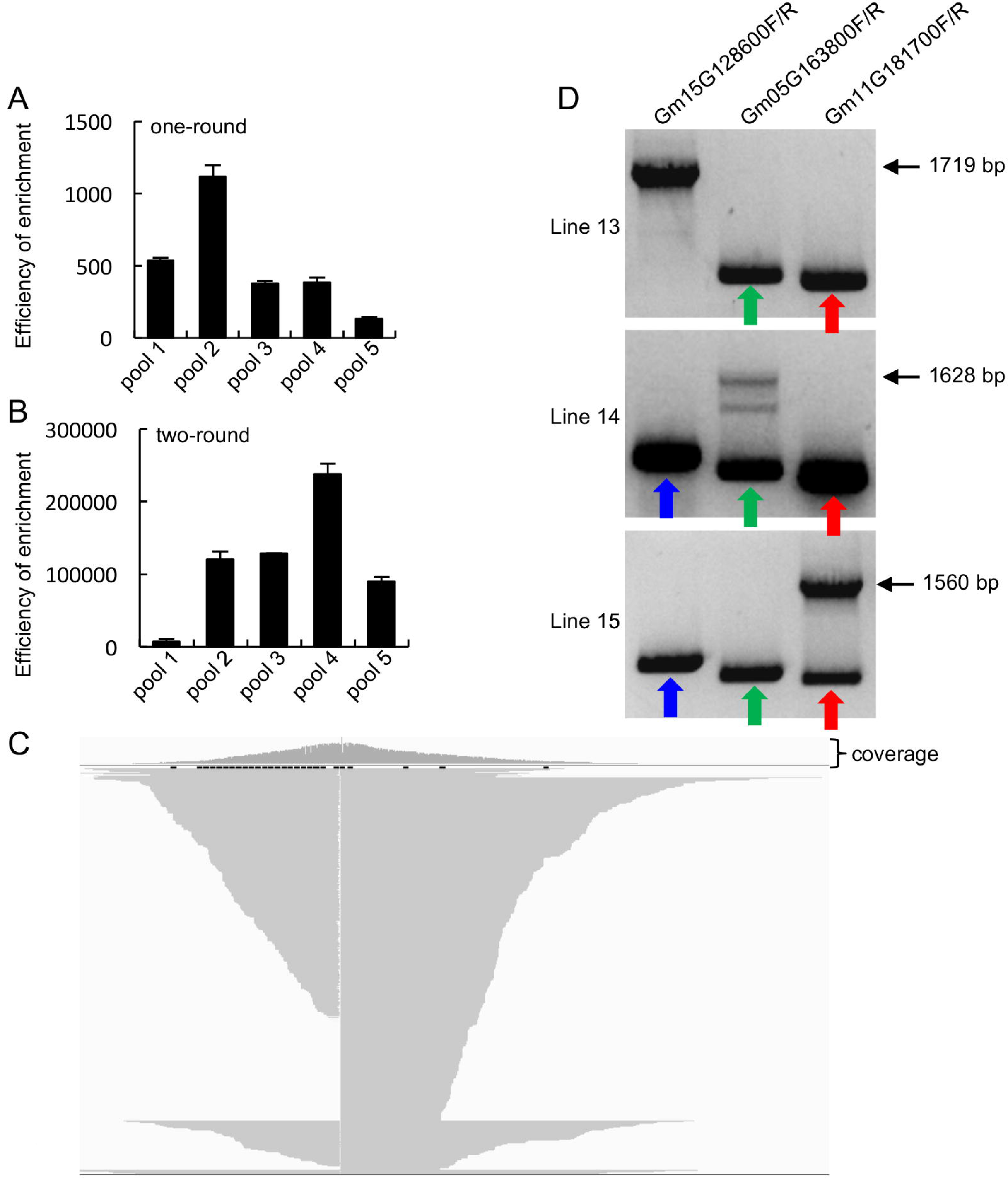
Sequencing results after two-round enrichment of the *Ds*-containing fragments. **(A)** and **(B)**Efficiency of one-round (A) and two-round (B) enrichment of the *Ds* element-containing fragments. 2% of samples before and after probe-enriching were used to perform qPCR. The amount of target fragments was normalized to that of internal control. **(C)**Schematic diagram of the flanking sequences of Line 15. Partial sequences of reads were shown. **(D)**PCR validation of the *Ds* insertion in Line 13, Line 14, and Line 15. The three individual lines were examined with three pairs of primers, respectively. Each primer pair (labeled above the picture) recognizes a potential insertion position of the *Ds* element, identified by sequencing. Line 13 containing a *Ds* insertion in Glyma15G128600 gene produced a 1719 bp fragment, while Line 14 and Line 15 without insertions in this gene generated ~ 559 bp fragments (indicated as arrows). Line 14 containing a *Ds* insertion in Glyma05G163800 gene produced a 1628 bp fragment, while Line 13 and Line 15 without insertions in this gene generated 462 bp fragments (indicated as arrows). Line 15 containing a *Ds* insertion in Glyma11G181700 gene produced a 1560 bp fragment, while Line 13 and Line 14 without insertions in this gene generated 394 bp fragments (indicated as arrows).

### MinION sequencing provides a platform for high-throughput method to identify map position of transgenic alleles in plants

Reads containing sequences of the target allele in soybean, a *Ds*-activation tag element, averaged in the hundreds (Table 3) from non-enriched genomic DNA, reflecting the power of MinION sequencing technology as a cost effective tool that could be translated as a high-throughput method to map a transgenic allele in the soybean genome. To further test its throughput, an expanded pooling was performed with the enrichment steps, wherein 51 independent soybean lines containing a single *Ds* element were divided into six pools, each of which contained eight to ten lines (Table 4), for minion sequencing. The outcome from this expanded throughput evaluation resulted in total read counts ranging from 19758 to 282690 across the pools, with reads containing the *Ds* sequence ranging from 212 to 16146 (Table 4). These data were sufficient to successfully map the transgenic allele in each of the 51 soybean lines analyzed (Supplementary Table S3).

**Table 4.**
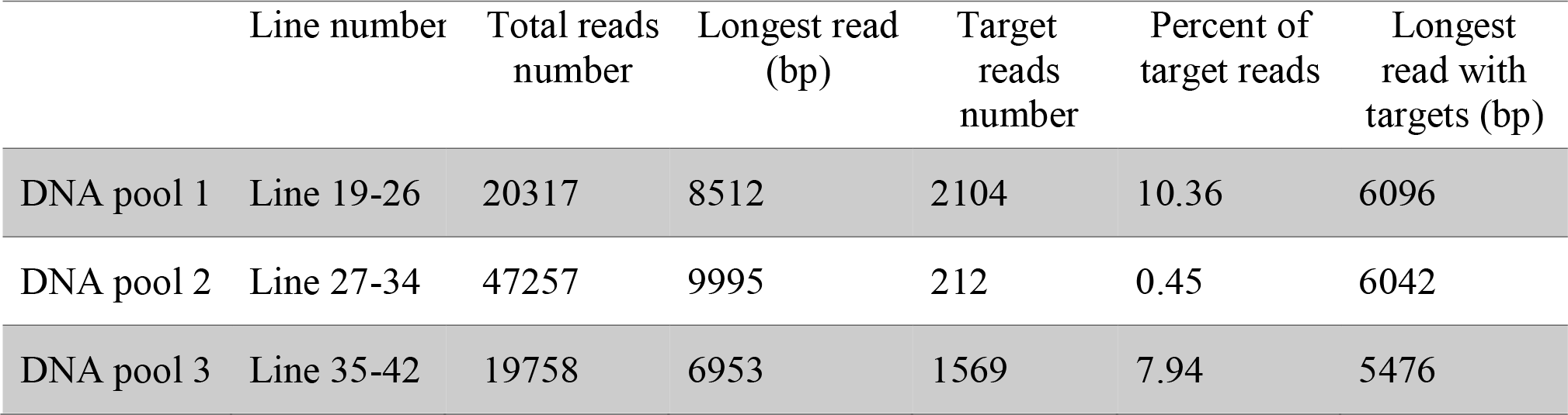
Sequencing result of the 50-sample pools.

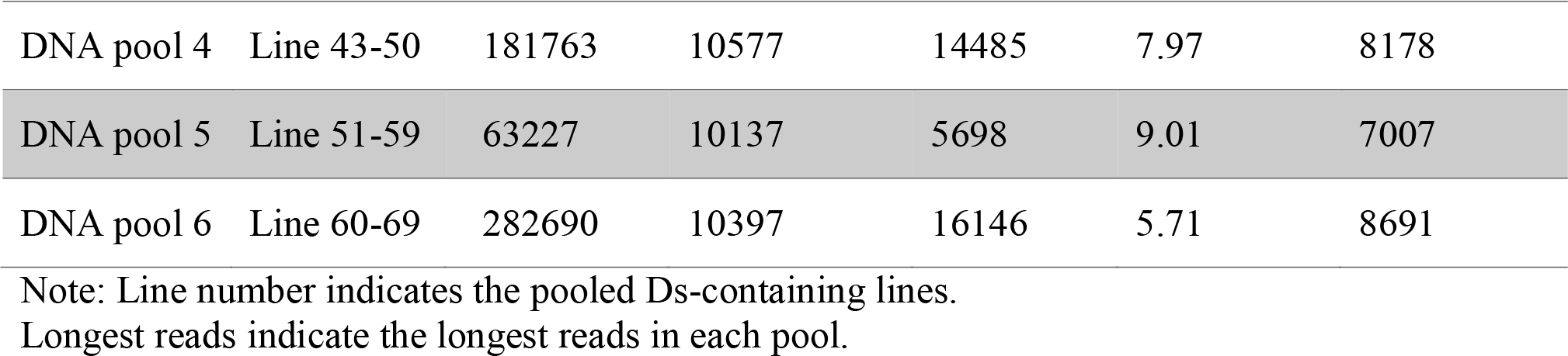

To further validate if this method is suitable to map potential multiple transgene insertions, we selected 18 transgenic soybean lines harboring one to 3 copies of the original Ds transgene (~ 5Kb), which were determined by southern blot (Figure S4 and Supplementary Table S4). We divided these plants into 4 pools, and performed MinIon sequencing after target enrichment. We were able to identify 29 transgenic insertion loci (Supplementary Table S3), which agree with the southern blot result. This result suggests that our method can be used to map transgene loci with known insertion numbers.

## Discussion

Communicated herein is a long read and affordable sequencing-method suitable for high-throughput mapping of transgenic alleles in higher plants. This method has at least five advantages. First, it provides reliable information of sequences flanking the insertion position. In most scenarios, over a hundred reads contain the target transgenic allele and associated junction sequences. Second, the method is scalable, by coupling pooling with enrichment steps prior to sequencing the transgenic allele in 51 independent lines were successfully mapped in a single sequencing run. Importantly, the reads containing the target allele are sufficient to accurately map a transgenic allele back to a reference genome. Thus, it is likely that sample pools can be further enlarged. Moreover, the target enrichment method still has potential for additional refinement given the ratios of reads containing the target sequences per total read count are still low (ranging from ~0.5 to 10%). The current enrichment step only incorporates one probe to the target allele. A refinement in the enrichment step might include the use of multiple probes that recognize different regions of the target allele thereby improving specificity and efficiency of capture. Third, the cost per map is relatively low, estimated at $1360 per 51 samples, excluding labor. If pooling can be expanded, the cost will be further reduced. In addition, after one round purification, we may also use primers that recognize the target and adaptor to amplify the target containing fragments, which will eliminate second round purification and improve specificity, and thereby reduce the cost and allow pooling more samples. Fourth, it is rapid with the timeframe from DNA fragmentation to mapped transgenic allele being approximately one week. Lastly, since MinION is a portable device that can run at a laptop or desktop computer, permitting utilization of this tool to modestly equipped laboratories globally, it has unrivaled convenience and broad usability.

The introduction of novel genetic variation into higher plants through the tools of transgenic technology offers a powerful way to complement plant breeding programs. Prior knowledge of transgene insertion position facilitates breeding decisions. The MinION-based sequencing strategy outlined here is a powerful, high-throughput tool to determine the insertion position of transgenic alleles in higher plants. The average length of reads containing the Ds element here was ~ 2.1 Kb and the longest reads was ~ 10 kb in the 51 sample-sequencing. This length should be sufficient to cover a portion of a longer transgene with flanking sequencing at one end. Indeed, we used this method to determine a population of soybean lines containing a ~5kb transgene. The average reads containing the Ds elements are more than one hundred, which should be sufficient to identify multiple insertion events in the genome. However, it may still be a challenge to identify transgene copy numbers with the current target enrichment method when multiple copies of the transgene exist in the same location of the genome. In this scenario, the average read length needs to be improved. A possible solution is to perform size selection after each round of target enrichment or after adapter addition to eliminate the short DNA fragments, and thereby to improve the read length, although this may reduce the numbers of reads containing the transgene.

Although this method is developed to examine a population of soybean lines containing the same transgene, it can be adapted to map the transgene insertion from plants containing different transgenes that do not share common fragments using probes targeting individual transgenes. We noticed variations of reading within a barcode. This may due to the difference of DNAs surrounding the insertion positions, which results in variations in efficiency of ligation or PCR. Moreover, there are reading variations among different pools. This is likely due to that different barcodes may have different optimal PCR conditions, as we currently use the same PCR amplification condition for all pools.

## Acknowledgments

This work was supported by the National Institute of Health (GM127414 to B.Y), the National Science Foundation (Awards OIA-1557417 to B.Y, C.Z, E.C and T.C, IOS-1444581 to T.C, and MCB-1808182 to B.Y), the Nebraska Soybean Board (Award #1727 to B.Y and #1728 to C.Z) and by a University of Nebraska, Agricultural Research Division grant to D.H.

## Supplementary data

**Fig. S1. The diagram of the *Ds* system.**

**Fig. S2. Agarose gel electrophoresis of sheared DNA. 8 ug genomic DNA samples in 150 ul ddH2O were fragmented to 8 kb**

**Fig. S3. Size distribution of readings containing the Ds elements.**

**Fig. S4. Copy numbers in various soybean transgenic lines determined by Southern Blot.**

**Table S1. Oligo DNAs used in this study**

**Table S2. Positions of *Ds* insertion identified in the 15-sample sequencing**

**Table S3. Positions of *Ds* insertion identified in the 50-sample sequencing**

**Table S4. Positions of *T-DNA* insertion identified in the 18-sample sequencing**

